# Pharmaceutical patent landscaping: A novel approach to understand patents from the drug discovery perspective

**DOI:** 10.1101/2023.02.10.527980

**Authors:** Yojana Gadiya, Philip Gribbon, Martin Hofmann-Apitius, Andrea Zaliani

## Abstract

Patents play a crucial role in the drug discovery process by providing legal protection for discoveries and incentivising investments in research and development. By identifying patterns within patent data resources, researchers can gain insight into the market trends and priorities of the pharmaceutical and biotechnology industries, as well as provide additional perspectives on more fundamental aspects such as the emergence of potential new drug targets. In this paper, we used the patent enrichment tool, PEMT, to extract, integrate, and analyse patent literature for rare diseases (RD) and Alzheimer’s disease (AD). This is followed by a systematic review of the underlying patent landscape to decipher trends and applications in patents for these diseases. To do so, we discuss prominent organisations involved in drug discovery research in AD and RD. This allows us to gain an understanding of the importance of AD and RD from specific organisational (pharmaceutical or university) perspectives. Next, we analyse the historical focus of patents in relation to individual therapeutic targets and correlate them with market scenarios allowing the identification of prominent targets for a disease. Lastly, we identified drug repurposing activities within the two diseases with the help of patents. This resulted in identifying existing repurposed drugs and novel potential therapeutic approaches applicable to the indication areas. The study demonstrates the expanded applicability of patent documents from legal to drug discovery, design, and research, thus, providing a valuable resource for future drug discovery efforts. Moreover, this study is an attempt towards understanding the importance of data underlying patent documents and raising the need for preparing the data for machine learning-based applications.

## 1. Introduction

Patent documents are considered crucial assets within the drug discovery domain as they allow the inventor rights over an invention for typically 20 years post filing. In the biomedical domain, these inventions could include information on drug formulation, dosage or efficacy, as well as information on the medicinal chemistry properties of leads or pre-clinical candidates. Moreover, the legal value of intellectual property in patent documents, and their distinctive function compared to scientific literature, makes patents crucial milestones in drug discovery and development [1]. Despite this, patent documents have been relatively untapped as a source to assist in scientific discovery and are used in the latter parts of drug discovery to assist investors in drafting new patents for their innovation or extension of existing patents in case of “me-too” drugs [2]. Also, the use of legal phraseology within patent documents requires experts in both patent drafting and patent analysis to evaluate and understand the underlying content. As a result, patent literature mining has developed into a specialised field that is closely linked with chemoinformatic analysis and commercial evaluation, meaning the appearance of patent sourced information is less common in publicly available scientific data resources. In the current context, we use the word “patent” to indicate both patent applications and granted patents.

Recently, there has been an increase in the attention given to patent documents as an aid to monitor advancements in drug discovery. This is seen in the field of oncology drug discovery, where patenting activity is intense, and patents offer a window into the latest techniques in translational cancer therapies [3–5]. To support these efforts, oncology-specific patent datasets have been catalogued by established patent offices such as the USPTO Cancer Moonshot Patent Data (https://www.uspto.gov/ip-policy/economic-research/research-datasets/cancer-moonshot-patent-data). Moreover, there have been efforts in mining and ingesting patent-related information into knowledge graphs. One such example is the Chinese patent medicine (CPM) knowledge graph, where a natural language model (BERT) was used to extract biological entities (specifically chemicals, diseases, and conditions) from patent documents [6]. The resultant graph assisted in the identification of disease gaps as well as summarising drug prescription trends in China.

In addition to the extraction of patent-related information, the importance of exploring patent literature has attracted new market interest from scientific information vendors. The value of an exhaustive patent exploration process is significant for the pharmaceutical industry, resulting in the establishment of commercial and open-source patent databases besides institutional offices such as the United States Patent and Trademark Office (USPTO), European Patent Office (EPO), and Japan Patent Office (JPO). The dominant commercial providers in this market are Clarivate (https://clarivate.com/products/ip-intelligence/ip-data-and-apis/derwent-world-patents-index/) and CAS-Scifinder (https://www.cas.org/solutions/cas-scifinder-discovery-platform/cas-scifinder). Simultaneously, public chemical databases have also incorporated patent information for their catalogued chemical collections. PubChem [7] has historically incorporated compound links to USPTO, while the European counterpart, ChEMBL, has developed a stand-alone patent database (SureChEMBL) [8] based upon annotated biomedical entities which allow for searching a larger chemical space to inform drug discovery. Within the drug development domain, SureChEMBL represents one of the largest open-access resources of patents with its ability to extract data from independent patent offices. Such databases have enabled researchers to systematically assess, identify, and explore patterns within patent documents [9, 10].

Patent documents can serve as a vast data source for machine learning (ML) and artificial intelligence (AI) applications [11]. ML-based algorithms can be used to predict the success rate of patented compounds by incorporating clinical trial results. This could be also beneficial for patent applicants in making R&D strategic decisions related to competitors. However, the unstructured nature of patent documents and their actual digital media, which are often in PDF format when downloaded from public repositories, also poses a challenge and as a result, manual curation is required to extract valuable information, such as bioactivity data, from these documents [12]. Therefore, there is an increasing need for tools for patent automatic analysis utilising language models (LM) to identify and extract relevant metadata for exhaustive patent data search.

In 2022, we introduced PEMT, a patent enrichment tool that allows extraction of pharmaceutically relevant entities such as targets and chemicals of interest from patent documents [13]. This tool annotates modulators with patent information rather than providing an overview of the usability of the patents in drug discovery. This manuscript aims to guide readers through the tool’s implementation and, in doing so, highlight the role patent literature can play in driving decisions in the drug discovery process. We utilised knowledge graphs (KGs), which are graphical databases composed of aggregated literature and experimental data, as the platform for demonstrating the utility of our analyses. For rare diseases, we utilised OrphaNet [14], while for Alzheimer’s disease, we used the Human Brain Pharmacome [15]. By means of these KGs, we extracted data from the past two decades of chemical patent space and performed patent landscaping, which involved the identification of patterns within patent documents. We start by identifying the commercial and non-commercial pharmaceutical originators based on the patent activity of their affiliated owners. Next, we performed a retrospective overview of targeted proteins from patent documents, to identify their importance in the drug development process at specific periods. Additionally, we examined the impact clinical trials have had on the corresponding target prioritisation. Lastly, we examined the drug repurposing aspect of pre-clinical small molecules and clinical stage drugs, highlighting potential novel disease treatment options. Overall, this patent landscaping approach highlights the importance of analysing patents and their usefulness in making decisions during drug development efforts.

## 2. Methods

In the initial sections, we discuss the KGs employed as data sources, explain the functioning of the PEMT tool, and outline the curation process necessary for standardising information from patent documents. Finally, we provide information on the implementation details pertinent to the analysis.

### 2.1. Knowledge graph selection

We extracted data from two indication-specific semantically organised KGs: OrphaNet and Human Brain Pharamcome (HBP). OrphaNet (accessed on 2021-11-01) is an open-source network that focuses on rare diseases and includes information on biological entities such as orphan gene targets and drugs, as well as clinical entities such as clinical trials and biobanks [14]. In contrast, the Human Brain Pharmacome (HBP) is a publicly available biomedical knowledge graph representing neurodegenerative diseases with a particular focus on Alzheimer’s disease [15]. The data in the graph is built on the Biological Expression Language (BEL) framework and includes information on proteins, biological processes, and other relevant features [16].

### 2.2. Experimental and patent data extraction

To expand the data resource with patent data, we used the PEMT tool [13], which systematically retrieves patent documents using a list of genes or proteins. To do so, it extracts manually annotated and verified chemical and biological modulators that have binding or functional effects on proteins from ChEMBL and subsequently captures all patent documents related to the modulators using SureChEMBL. The patent documents collected are then filtered based on their pharmaceutical relevance and status (i.e. whether the patents are active or expired). Thus, the tool removed patent documents older than 20 years and only included those documents that were tagged with specific IPC code classes referenced in the publication [13]. These codes are alphanumeric codes assigned by officials to patent documents and help in understanding the scope of the patent.

### 2.3. Patent data harmonisation pipeline

To aid systematic analysis, we developed data harmonisation and normalisation pipelines. In this pipeline, patent owner names were classified into three categories: “Organisation”, “Acquired”, and “Individuals”. Organisations include patent owners that were part of companies and universities, while Acquired owners are those that were within legacy entities which have been acquired or merged into larger Organisations. An example would be Dupont Pharmaceuticals, which Bristol Myers Squibb (BMS) acquired. Individuals are individuals who own a patent. Additionally, similar organisations were grouped for consistency. For example, all variations of Bristol Myers Squibb were mapped to Bristol Myers Squibb. This normalisation step allows for identifying all relevant patents from the same owner and has not been performed previously.

### 2.4. Compound clinical stage annotation

To annotate compounds based on their clinical trial stage, we utilised the information captured within ChEMBL. For each compound registered in ChEMBL, an additional annotation is made based on whether the compound has been a clinical candidate and if so, at which stage. As a result, ChEMBL assigns phase numbers 0 - 4 where 0 indicates a compound being researched in the preclinical stage, 1 to 3 indicate the compound being tested in clinical trial phases I - III, and 4 indicates the compound has been approved by FDA and/or EMA. In our analysis, we make use of this ChEMBL data to identify compounds, with associated patents, that have been moved from the preclinical to the clinical setup within the drug discovery pipeline.

### 2.5. Patent data clustering

To cluster patent owners, we made use of a hierarchical approach where two or more owners are clustered together based on a distance metric from singletons to cluster groups in a bottom-to-top manner. This type of clustering allows creation of dendrograms that help establish patent owner groups that belong to the same cluster. The distance metric used in the clustering approach was the distance correlation which enabled in measuring the dependency between the two owners **(Equation 1)**. The clustering was achieved using Seaborn ‘clustermap’ package (https://seaborn.pydata.org/generated/seaborn.clustermap.html).

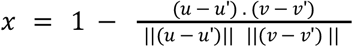

**Equation 1. Distance correlation.** The distance correlation is a metric used to clusters that have minimal distance together. In this equation, *u*′ is the mean of vector *u* and *x . y* is the dot product of *x* and *y*.

### 2.6. Target based publication data collection

To extract publications relevant to targets of interest, we made use of open source platforms within NCBI Gene Browser (https://www.ncbi.nlm.nih.gov/gene/) to gather information on the human targets. Through the link between the gene platform and PubMed (https://pubmed.ncbi.nlm.nih.gov/), we collected all relevant publications related to a target. For our purposes, we clustered all the publication found in a year together and used this statistics as the basis for understanding the research trend on targets. Moreover, we rescaled the publication counts to be between 0 and 1 using the min-max normaliser (**Equation 2**).

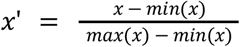

**Equation 2. Min-max normalizer.** The min-max normalise allows us to scale the values to a define range. In this equation, *x* is an original value and *x’* is the normalized value.

### 2.7. Data availability and implementation details

To run our PEMT tool, we extracted human proteins that modulate a disease or symptom from both graphs. The PEMT tool was run on a Windows operating system and was followed by the harmonisation pipeline mentioned in **Section 2.3**. Additionally, within the collected patent documents, we found 131 patents in rare diseases and 705 patents in Alzheimer’s diseases with no associated owner. This is likely due to the quarterly updation of public databases like SureChEMBL, causing a lag in the time for updating. As a result, additional filtering was done to exclude patents with no owners from the study. **Figure 1** provides an overview of the process involved in collecting and analysing the patent dataset, referred to as the patent corpora, in the following sections. The data and the scripts used during this analysis can be found on GitHub at https://github.com/Fraunhofer-ITMP/Pharmaceutical-patent-landscaping.

**Figure 1.**
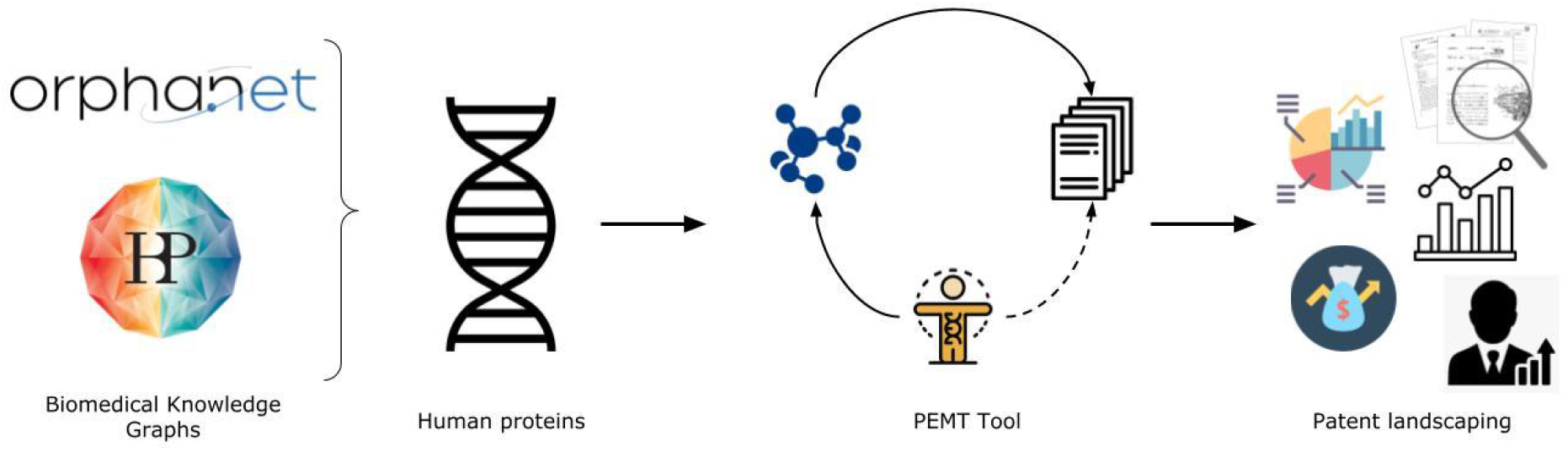
Synopsis of the patenting landscape analysis. We collect all human proteins from the two biological networks relevant to diseases and symptoms. Using these proteins as a starting point, we run our PEMT tool and retrieve all associated modulators and patent documents based on predefined conditions. Finally, using the patent document dataset, also referred to as patent corpora, we conduct deductive and exploratory analysis to uncover patterns within the dataset.

## 3. Results

We analysed the patent extracted from 4,314 and 10,237 proteins from OrphaNet and HBP respectively. In this analysis, we define “rare diseases” as diseases mentioned within the Orphanet database. In the following section, details four subsections through which we explain how patent literature can be utilised in drug discovery. First, we provide an overview of the information collected from the patent corpora for each disease. Next, we retrospectively analyse the patent corpora unveiling leading organisations in drug research and discovery (R&D) for individual diseases. Following this, we investigated case studies for selected targets based on biological networks and examined their patenting trend. Lastly, we illustrate the usefulness of patent documents in identifying drug repurposing opportunities.

### 3.1. Overview of the patent corpora

In this section, we investigate the patent corpora of each disease and provide a statistical overview of the information collected within the patent corpora. This summary will provide insights into the scope of patenting activity within the disease area, denoting whether the patents are filed for pharmacological relevance or for formulation purposes. We will also examine and compare the number of patent documents that have been approved versus those still in the application pipeline.

We retrieved 17,506 compound-patent pairs from the dataset, which included 585 compounds and 502 unique patent documents with the rare disease patent corpora. Out of the patents retrieved, 180 were granted patents, represented by documents with kind codes starting with B or E, while the rest were still in the application process, represented by documents with kind codes starting with A (**Figure 2A)**. Additionally, based on the IPC classification, it was revealed that more than half of the patents were related to C07D (https://www.wipo.int/classifications/ipc/en/ITsupport/Version20170101/transformations/ipc/20170101/en/htm/C07D.htm#C07D), which includes heterocyclic compounds. Following this, 12% of patent documents were in the IPC code C07J (steroids), 7% were in code C07K (peptides), and 6% were in code A61P (specific therapeutic activity of chemical compounds or medicinal preparations) (**Figure 2B**).

**Figure 2.**
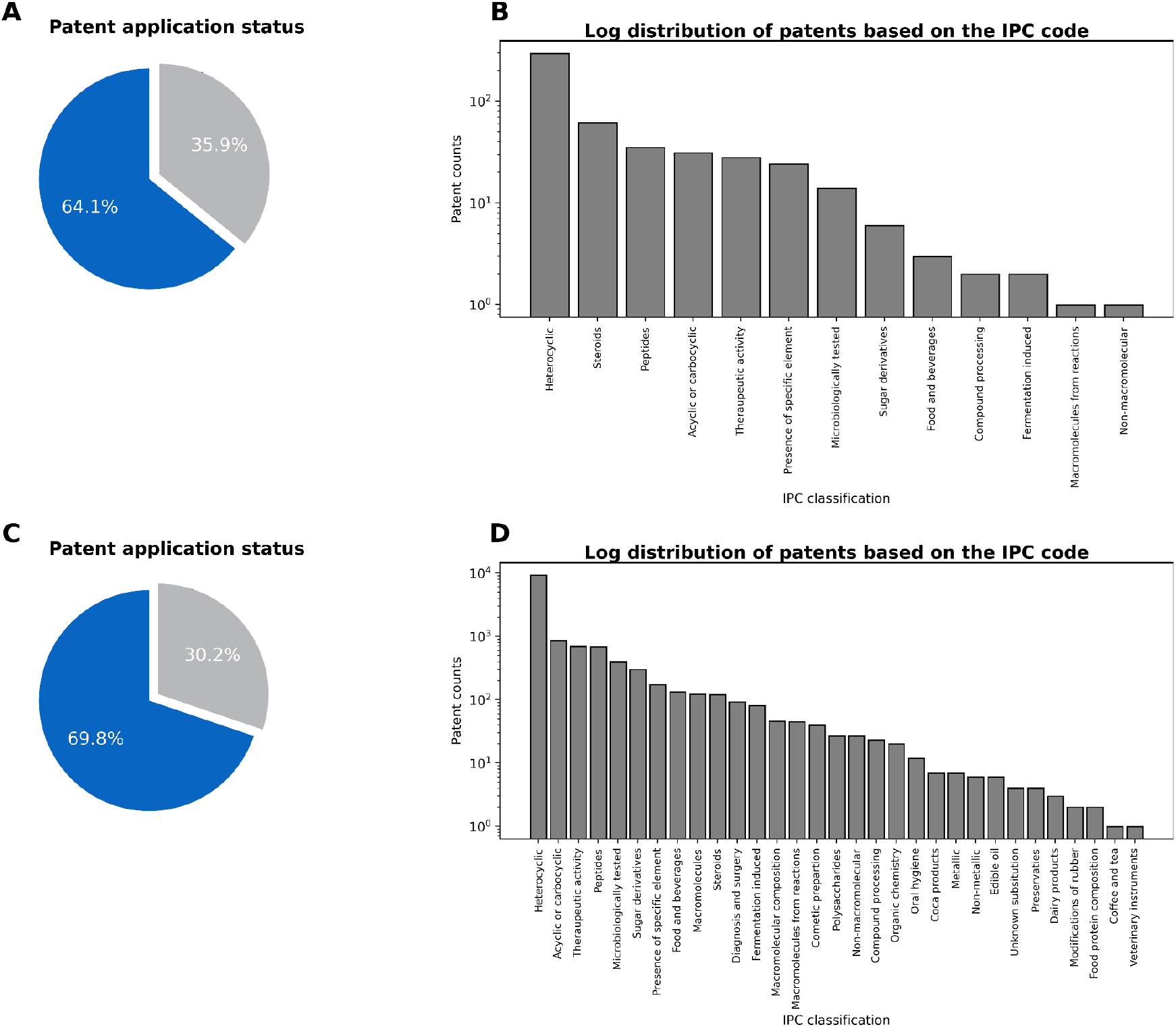
Patenting Landscape overview for Rare and Alzheimer’s Diseases. **A)** Rare disease patent application ratio denoting whether the patents have been granted (in grey) or are still pending (in blue). **B)** Logarithmic distribution of the rare disease patents based on the IPC classification category. **C)** Alzheimer’s disease patent application ratio denoting whether the patents have been granted (in grey) or are still pending (in blue). **D)** Logarithmic distribution of the Alzheimer’s disease patents based on the IPC classification category.

Conversely, in the case of the Alzheimer’s disease patent corpora, we retrieved 76,321 compound-patent pairs that included 22,930 compounds and 13,181 unique patent applications. From these patents, 3,980 were granted (indicated by kind codes starting with either B or E), while the remaining were still under the application process (indicated by kind codes starting with A) **(Figure 2C)**. Upon reviewing the IPC classification of the patents, it was found that 70% of the documents belonged to class C07D, representing patent applications mentioning heterocyclic compounds. Alongside this class, other notable IPCs include C07C (representing acyclic and carboxylic compounds) with 7% of patent documents, A61P (representing a specific therapeutic activity of chemical compounds or medicinal preparations) with 5% of patent applications, and C07K (representing peptides) with 5% of patent applications **(Figure 2D)**.

Thus, RD and AD patent cohorts revealed two recurring patterns: i) a lower number of granted patents and ii) a dominance of “C07D coded” patent applications. The observation that granted patents are approximately half of all applications filed is consistent with the statistics within the pharmaceutical domain generated by EPO (https://new.epo.org/en/statistics-centre#/technologyfields?code=16). This filed-to-granted patents ratio can be attributed to several reasons. One of the reasons is the difficulty and expenditure involved in the research and development around patents. With the limited number of individuals reviewing the documents, the search for potential competitiveness within pre-existing inventions is time-consuming (https://www.epo.org/learning/materials/inventors-handbook/protection/patents.html). Additionally, the prevalence of C07D patents is likely due to the diversity provided by this class of compounds, which makes them a desirable candidate for drug discovery patent innovation compared to other compound classes [17].

### 3.2. Trends in the patenting landscape from the patent owners’ perspective

Next, we analysed the patenting activity for each disease to determine the major pharmaceutical companies and academic institutions that have made significant contributions to the R&D of potential drugs in the field. To do so, we leveraged the high-level patent owner classification (i.e., organisation, acquired, and individuals) and grouped patent owners based on the number of applications they have filed. Then, we reviewed the top patent owners and deduced their historic patenting activity in the field. We examined RD and AD separately to understand better the patent landscaping related to each disease.

#### 3.2.1. Identifying key players in rare disease patenting over the past 20 years

For RD’s, from 2000 to 2021, 369 organisations, 53 individuals, and 80 acquired organisations have played a role in the patent landscape. As illustrated in **Figure 3**, patenting activity has grown over the years, with organisations being the most active, followed by individuals. The boost in the patenting of large organisations, mainly pharmaceutical companies, is ascribed to legislations like the Orphan Drug Act (ODA), introduced in 1983, which gave financial incentives for developing therapies for rare diseases [18]. On top of that, the surging patent activity within the field after 2012 was correlated to the establishment of the Patient Protection and Affordable Care Act (ACA) which catalysed engagement of investors in financing R&D for orphan drugs and diseases [19]. On the contrary, a decrease in independent patent applications by individuals was observed since its peak in 2013. This effect could be due to two reasons: firstly, changes made in US and EU legislatures during the early 2000s, moving the “professor’s privileges” at IP generation to the universities, and secondly, professors and professionals, due to their legislation, were either hired by commercial companies or became part of larger academic groups [20].

**Figure 3.**
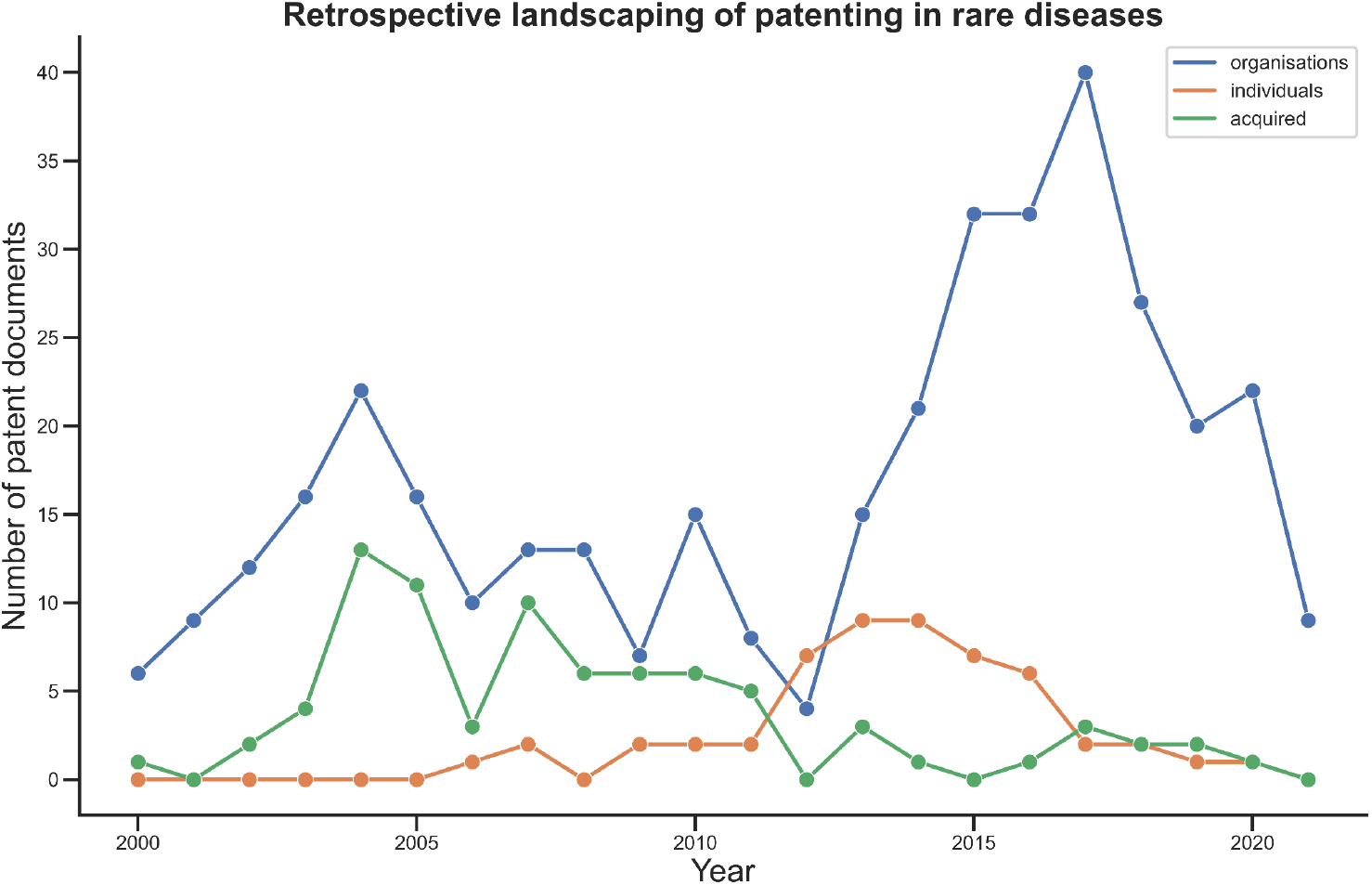
The retrospective landscape of the rare diseases patent corpora in the past two decades. The line plots indicate the growth of the patent documents years based on assignee type: organisations (blue), individuals (orange), and acquired (green). As expected, a large contribution is evident for organisations compared to others. A large boost in the Individual owners is seen around 2012, after which there has been a decline due to the legislatures.

To get a historical perspective of these patents, we selected 10 top patent owners and organised their patent applications based upon the timestamp of each patent application date, also known as the priority date. This subset of owners consisted only of global pharmaceutical companies such as Pfizer, GlaxoSmithKline, etc (**Figure 4**). It is essential to note that many start-up and medium-sized companies have emerged and subsequently been absorbed by larger organisations providing them with the leverage of patent acquisition filed by the smaller companies. One example is Sterix Ltd., which had 33 patents and was acquired by Ipsen (in 2004), adding to the company’s patent portfolio. Furthermore, most top assignees had a decreasing patenting activity in the past years. For instance, global giant Takeda, a Japanese multinational pharmaceutical company, just had one compound, Luvadaxistat, out of 122 from the patent compound space between 2000-2021 that went into the clinical trial phase II for Friedreich’s ataxia [21]. Thus, indicating a reason for decreasing interest of Takeda in rare diseases patenting **(Figure 4)**. Regardless, the top 10 assignees accounted for 44% of the 502 patent applications analysed, and the top 20 assignees accounted for 61%. Additionally, we observed the cumulative positive patenting trend for the top patent inventors in rare disease. The trend between 2013 and 2021, however, can be observed with a peak around 2016 and a gradual decline in the later years (**Supplementary Figure 1**).

**Figure 4.**
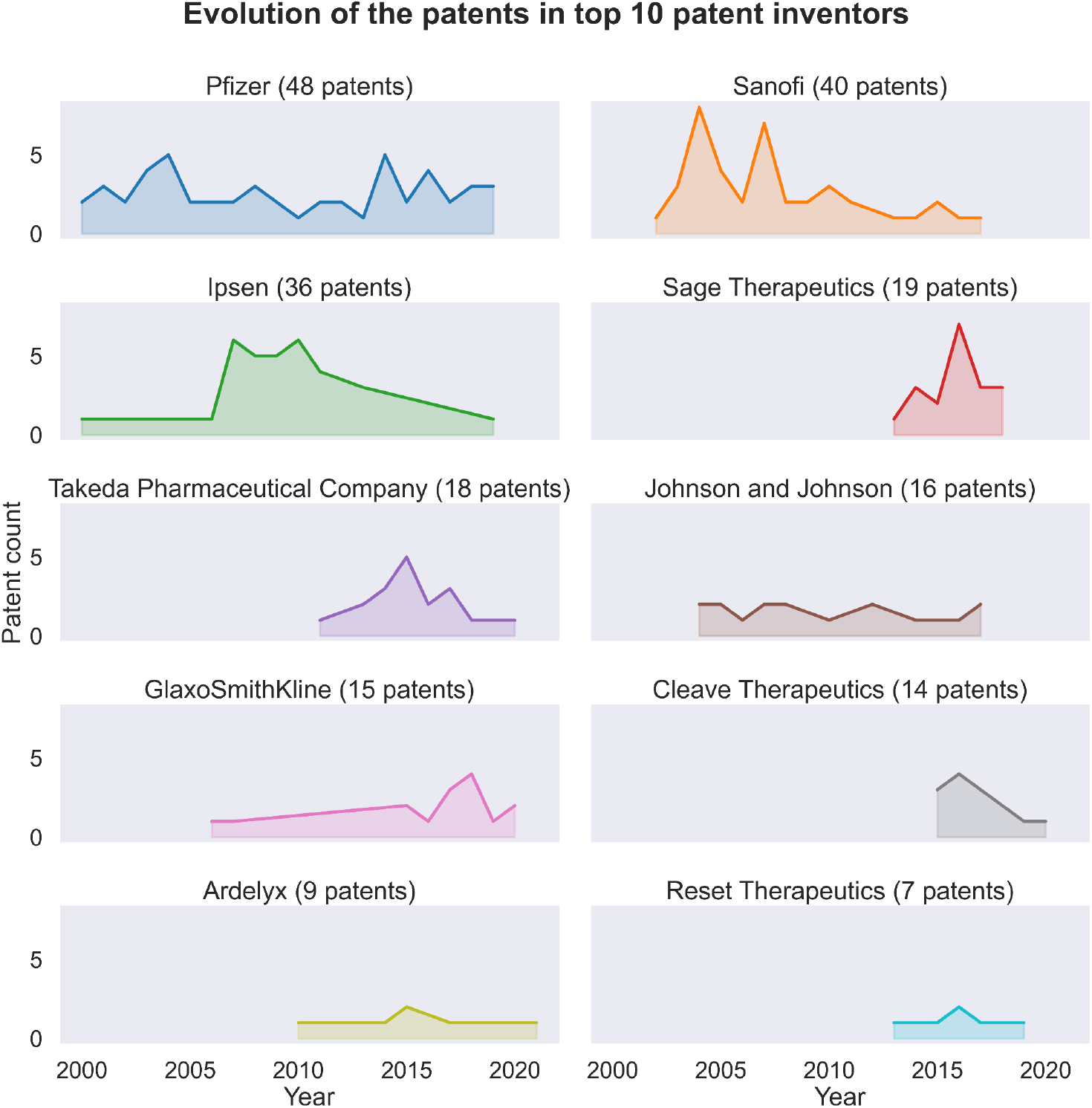
Top owners-based patent analysis in the field of rare diseases. Patent landscaping of the top 10 owners in the rare disease domain in descending order. The top owners were deduced from the In the figure, it can be seen that large pharmaceutical companies historically active in rare diseases, like Sanofi or Takeda, declined their patenting activity in the past few years, allowing other biopharmaceuticals such as GlaxoSmithKline, Pfizer, and Sage therapeutics to enter the field.

Additionally, we looked into the correlation between the patenting activity for the top owners based on the target portfolio to identify pharmaceutical companies that show similar target portfolios. We identified two clusters when correlating the target-based portfolio of the top 10 owners in rare diseases **(Figure 5)**. One of the clusters was between Sanofi and Ardelyx. We suspect this was due to their collaboration on sodium and potassium channel inhibitors (which include KCNK3, KCNK9 and SLC9A3), allowing them to apply for over 50 patents (https://ir.ardelyx.com/news-releases/news-release-details/ardelyx-licenses-nap2b-phosphate-inhibitor-program-kidney). A prominent target in the cluster space is DAO, with competitive patenting between large pharmaceutical companies like Pfizer, Takeda, and John and Johnson. A number of singletons are also seen, indicating the development of candidates targeting isolated targets (e.g GSK, Cleave). Together this target portfolio approach allows for identifying top patented targets within the indication areas and comparing patent owners’ target portfolios.

**Figure 5.**
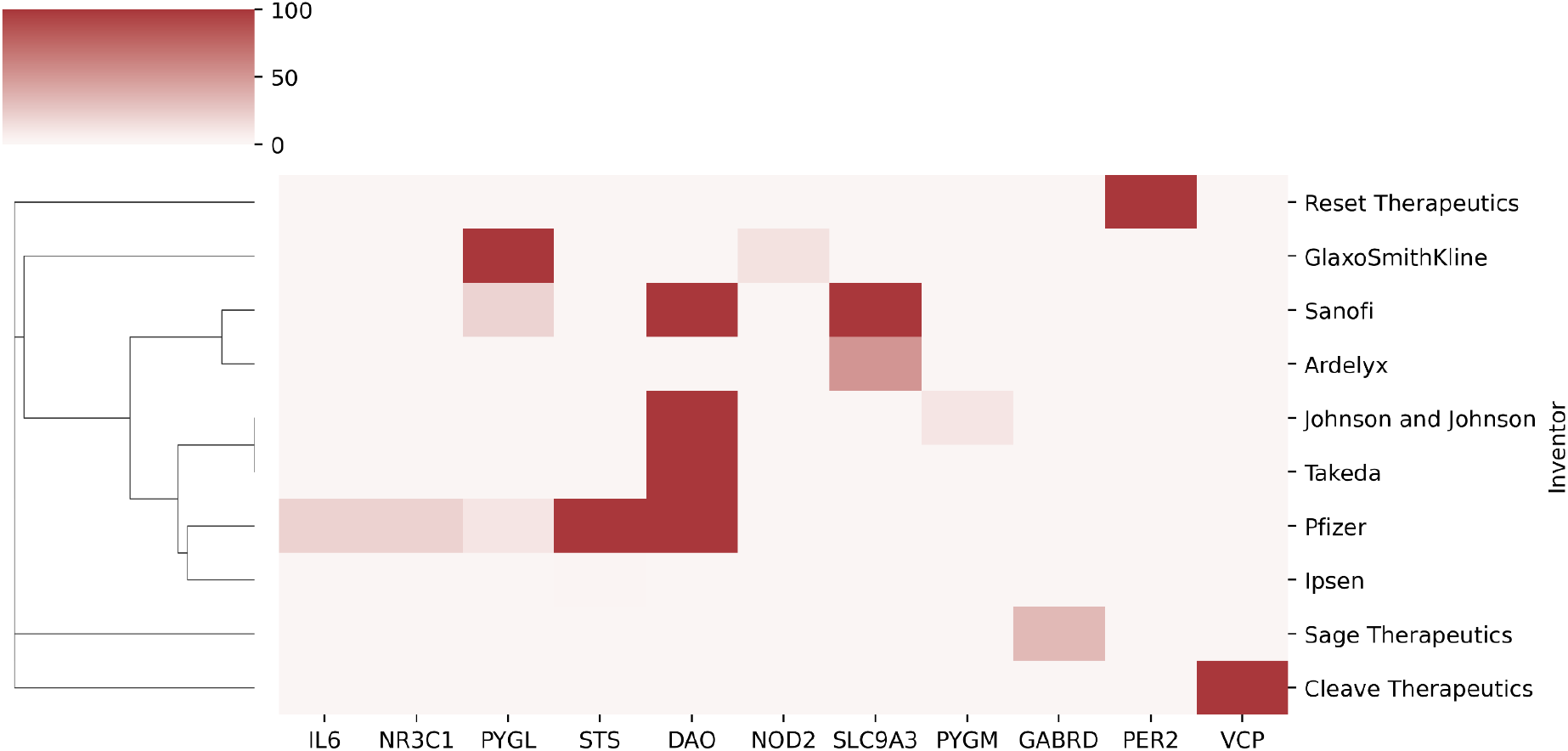
Target portfolio of top 10 owners in rare diseases. The hierarchically-clustered heatmap denotes the correlation distance between the top 10 rare disease owners based on their respective target patent landscape. The colour scale in the heatmap represents the number of patent documents associated with these targets. Together this clustering approach allows for the identification of clusters, here between Sanofi and Ardelyx and Pfizer and Ipsen, thereby indicating overlap between the targeting portfolio of these pharmaceuticals. From the plot, we can also notice the focus on DAO inhibitors within the rare disease industry.

As mentioned, no patent assignees in the top 10 were affiliated with an academic institute. To depict the difference, we divided organisations into two categories, industry and academia. Interestingly, a boost in patent applications from the academic sector was seen after 2012 has been tracked. (**Supplementary Figure 2A**). However, a general positive trend in patent applications by the industry is observed denoting increasing interest within pharmaceutical drug discovery for novel treatments for this disease. During the same time, patent law amendments took place in the United States of America, commonly referred to as the 2011 Patent Reform Act [22]. The reformed law pointed out the change in the prosecution of patent applications within the U.S. Patent and Trademark Office (USPTO) by moving from a “first-to-invent” to a “first-to-file” system. This law thereby redefined what was required in terms of prior art, modifying the application process in a significant way.

#### 3.2.2. Identifying key players in Alzheimer’s disease patenting over the past 20 years

In Alzheimer’s disease, 10,017 organisations, 1,778 individuals, and 1,386 acquired organisations have contributed towards the patenting activity. Similar to the rare disease patent landscape, organisations are in the lead compared to individuals and acquired organisations (as shown in **Figure 6**). This is attributed to the exceptionally high cost involved in the research and development of clinical trials, which cannot be afforded by publicly funded universities or small-scale organisations unless in partnerships with pharmaceuticals [23]. On the other hand, individuals showed an increase in patenting activity in 2011, with a peak in 2013 and then a decline afterwards, likely due to changes in patenting laws made by US and EU government bodies that allowed for commercial innovation rights to universities as well. Thus, individuals affiliated with universities lost their right to file patents to universities, while other individuals attracted pharmaceutical companies towards them and soon were hired by these large companies.

**Figure 6.**
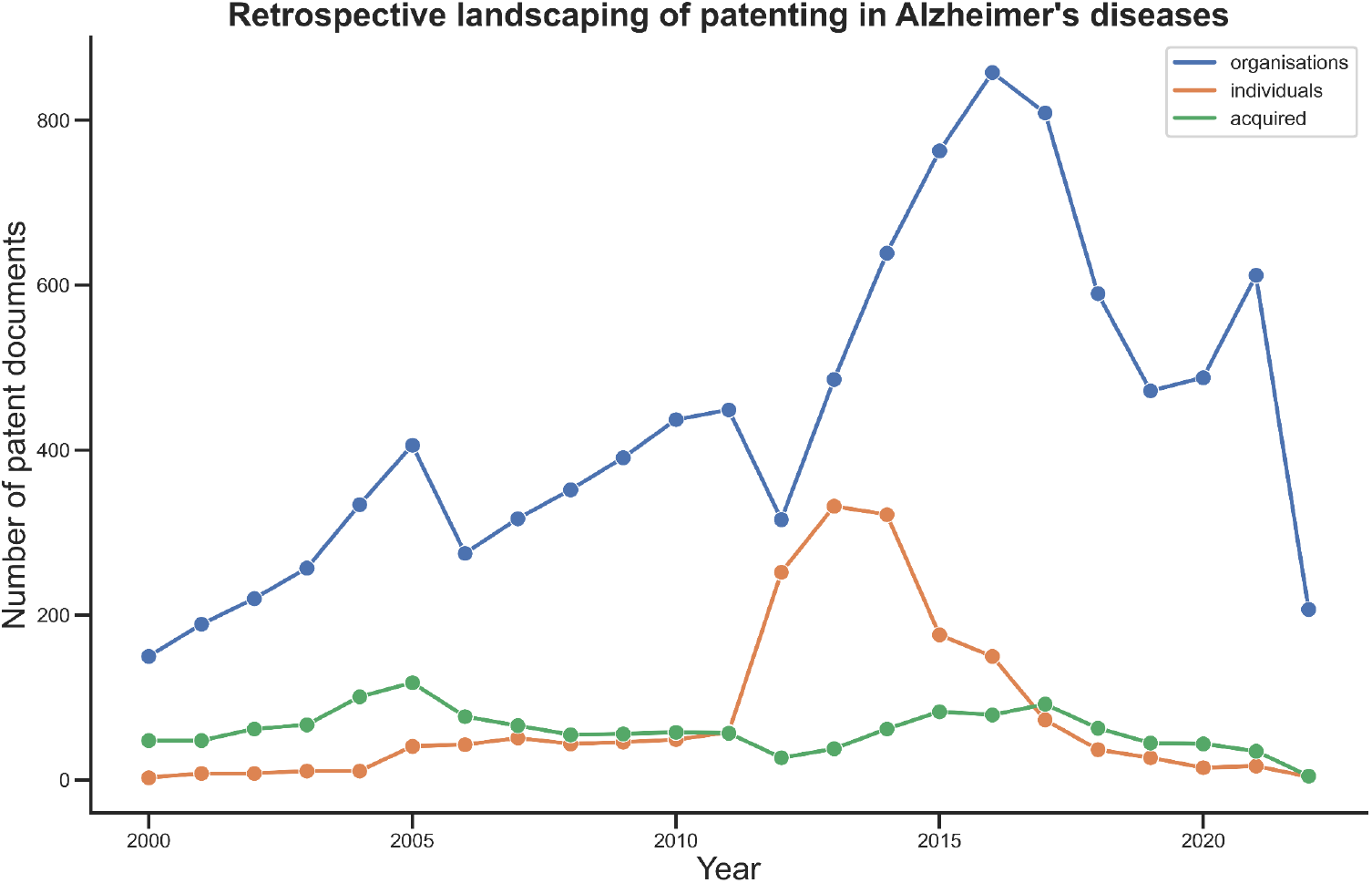
Patent activity by owner type in the Alzheimer’s field. The line plots indicate the growth of the patent documents years based on assignee type: organisations (blue), individuals (orange), and acquired (green). Pharmaceutical companies and universities are leading in patenting activity compared to others. Compared to the rare disease field, the contribution of the acquired owners is greater. A decline in Individual owner activity was observed post-2012 due to the legislature changes in US and EU.

We generated a retrospective perspective of the top 10 organisations found in Alzheimer’s patent corpora (**Figure 7**). Even though the total number of patent applications was higher for Alzheimer’s disease compared to rare diseases, a similar trend was observed. The top patent owners include Roche, Bristol Myers Squibb, Pfizer, etc., which are global pharmaceutical companies. There were “patent cliffs”, where a sudden loss of patent rights occurs due to concomitant expirations of several patents at the same time [24], within the underlying cohorts with decrements and increments in the patenting activity during the two decades. The increment denoted the increased interest of researchers in understanding the complex mechanism involved in Alzheimer’s disease. Despite the high investment in basic and clinical research, the mechanisms involved in the modulation of Alzheimer’s disease remained unclear, with a high attrition rate seen in clinical trials [25, 26]. We would like to point out that the trend mentioned above for Alzheimer’s disease would not be fully concordant with the current market trends as we did not consider any “biologicals” compounds, which have played an enormous role in drug discovery against Alzheimer’s disease lately [27]. A similar trend as that of rare diseases was observed in the cumulative patenting of the top patent inventors in Alzheimer’s disease. A gradual increase in activity was observed from 2013 with a peak around 2016 followed by a decline in the activity as the years came closer to the COVID-19 pandemic age (**Supplementary Figure 1**).

**Figure 7.**
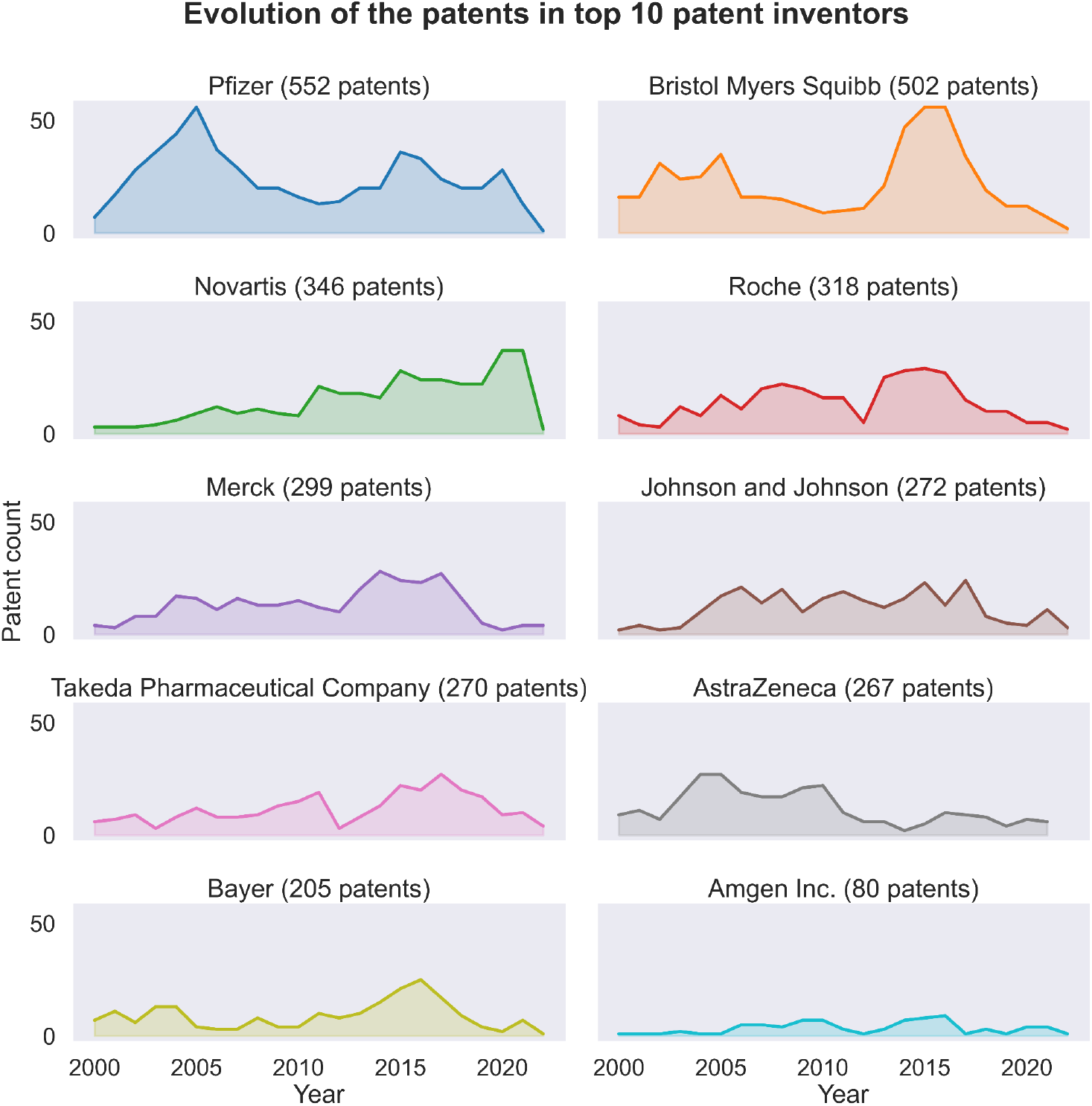
Patenting activity from the top owners in Alzheimer’s disease. In descending order, the plots showcase the patenting activity of the past two decades for the 10 top owners. The leaders in the field are global pharmaceutical companies such as Pfizer and BMS. All the top owners have been active within the field of Alzheimer’s with consistent patenting activity.

Next, we looked at the top patent owners with Alzheimer’s disease and plotted their target portfolio to examine whether two or more owners follow a similar target portfolio. Prominent collaboration partners such as Novartis and Amgen have coordinated neuroscience efforts and hence can be seen within a cluster in **Figure 8** (https://www.genengnews.com/news/amgen-novartis-launch-neuroscience-drug-collaboration/). Moreover, individual companies such as Takeda are observed due to their dominance, in particular, targeting mechanisms such as LPAR5 and SLC6A9. It is noteworthy that despite having common target spaces with leading pharmaceuticals, each top owner has at least one other neurodegenerative-related target space where it is leading with respect to the patenting landscape. ROCK2 for BMS, TTK for Bayer, and GCG for Pfizer are examples of this case. Overall, this approach enables identification of patent owners with comparable target focus and uncovers patterns in neurodegenerative drug target prioritisation.

**Figure 8.**
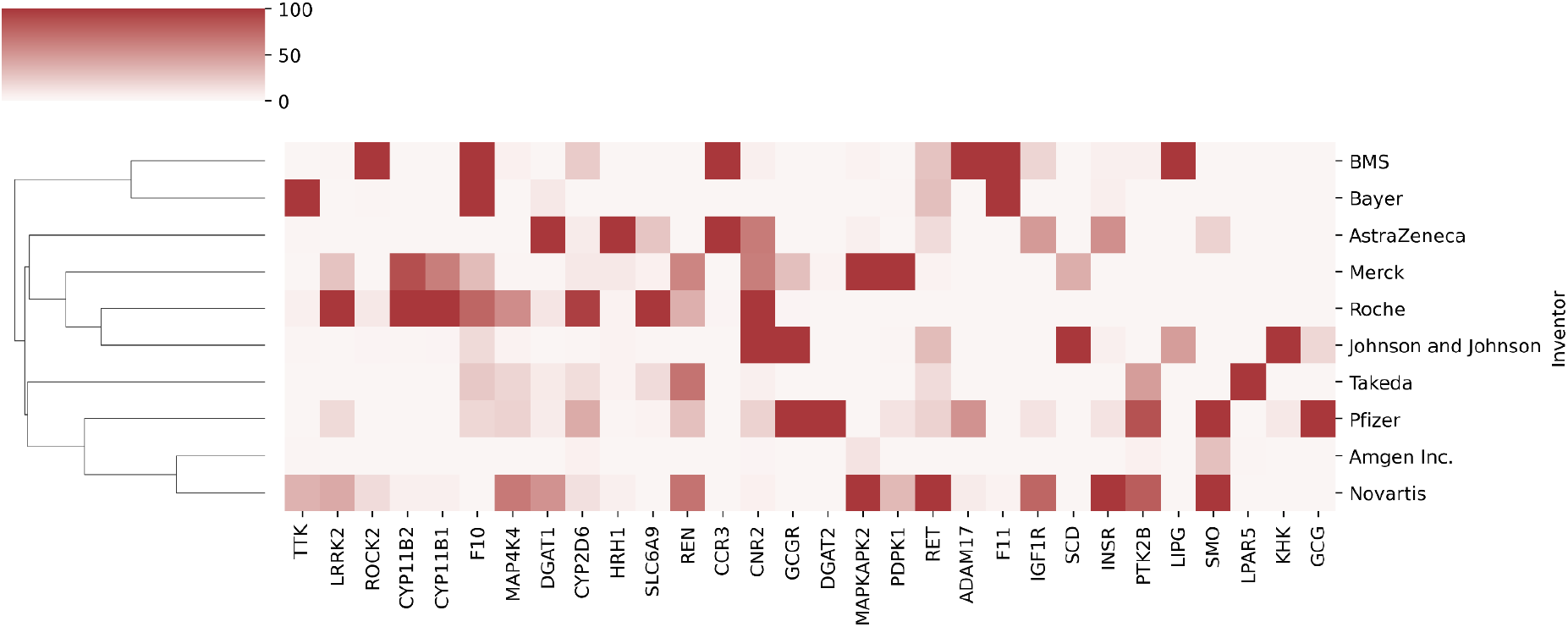
Target portfolio of top 10 patent owners in Alzheimer’s diseases. The hierarchically-clustered heatmap denotes the correlation distance between the top 10 Alzheimer’s patent owners based on their respective target patent landscape. Each coloured box in the plot indicates the number of patents from the owner linked to the specific target. We can observe three main clusters in this plot: the first is that of BMS and Bayer, the second is that of Roche and Johnson & Johnson, and the third is composed of Amgen and Novartis.

Similarly to the rare disease case, none of the top patent owners were from the academic sector, which is not surprising given the R&D cost involved in the case of Alzheimer’s disease [23]. Thus, we looked into the patenting activity of the academic sector and compared their growth with the industry (**Supplementary Figure 2B**). Similar to rare diseases, a general positive trend in patent applications by the industry is observed. An incremental rise in academic patenting activity was observed due to the amendment of the 2011 Patent Reform Act that was highlighted previously. During the same period, there was establishment of an act in the United States, the National Alzheimer’s Project Act (NAPA), that promoted research and development in the field of Alzheimer’s (https://www.nia.nih.gov/about/nia-and-national-plan-address-alzheimers-disease). In Europe, collaborative projects involving public-private partnerships with key stakeholders from the pharmaceutical industry were developed. One such project was the European Prevention of Alzheimer’s Dementia Consortium EPAD (https://ep-ad.org/) which involved data collection and analysis to speed up the drug discovery process for the disease, and support for improving clinical trial practice and design.

### 3.3. Elucidation of the research trends of targets using patent documents

Many factors contribute to the successful discovery of small molecule drugs for specific therapies. One of the essential factors is the mode of action (MoA) of the drug in the human body, which refers to the effect a drug has on a biological protein (also known as a target) that is relevant to mechanisms involved within diseased conditions. With the help of patent documents, we can investigate the target-based MoA claims within patent applications based on their effect in certain indication areas or on their importance over time from success achieved in clinical trials. Additionally, identifying potential disease targets, their prevalence, and their role in affected patient populations can be useful in developing new intellectual property. In this section, we will focus only on trends in the prioritisation of targets in patent literature over the years, while the therapeutic usage of targets has been previously described [28].

To investigate the research trend for targets, we first linked the patent information to targets. This was done by retrospectively connecting the patents to the targets through small molecule modulators. We would like to highlight that no prospective mapping of patents to targets based on their presence in the patent document was performed to confirm the link. Similar to the patent assignee analysis, we looked into the prevalence of targets over time **(Supplementary Tables 1 and 2)**. Furthermore, we will clearly distinguish the target priorities between RD and AD to understand the involved targets’ evolution. For both RD’s and AD, only the top targets, represented by the number of corresponding patent documents, were used to understand the target portfolio focus within the pharmaceutical companies.

We first started to evaluate the variations of the targets that are prioritised each year, to form the basis for target-centric landscaping, enabling us to identify targets that have been continued, diverted (to other targets), or halted, with respect to research, during their development. During this analysis, it was observed that the target perspective of the patent landscape was characterised by waves, which meant that not all targets were prioritised simultaneously **(Supplementary Figure 3)**. Thus, we looked at selected high profile targets for each disease and compared the patent activity with the corresponding therapeutical trial activities to understand the correlation between the agglomerated targets in a specific year and the preclinical research in drug discovery.

Glycogen phosphorylase L (PYGL) was one of the high incidence targets with 71 patent references. The mutation in the PYGL gene causes inhibition in the conversion of glycogen to glucose, thus, associating it with an autosomal recessive rare disease called glycogen storage disease type VI (GSDVI) [29]. This key link between the mutant PYGL and GSDVI was the reason for its early research and patenting activity. However, over the past years, interest in this target has gradually decreased, with almost no patent documents between 2014-2019 **(Figure 9A)**. In early 2019, small molecules called glucokinase activators (GKAs) were thought to be a potential solution for type II diabetes due to their ability to regulate glucose-6-phosphate and PYGL [30]. As research progressed on GKAs, it was found that they induced hypoglycemia, a condition characterised by decreased blood glucose levels, causing all clinical trials during that time to be terminated [31]. As a result, researchers refocused on the PYGL gene post-2014.

**Figure 9.**
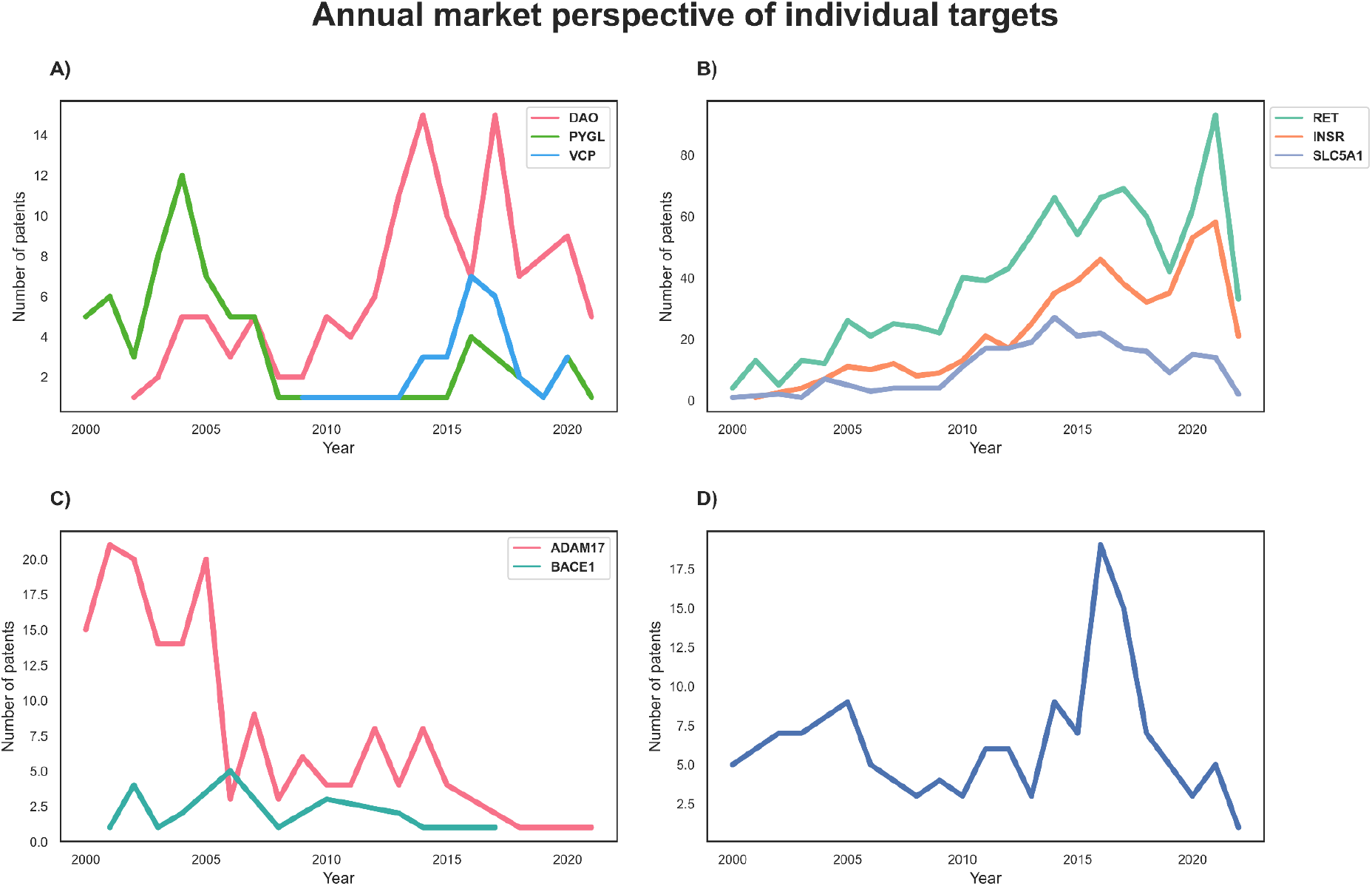
The patenting frequency of selected targets within each disease domain. **A) Targets for rare diseases**. In the case of rare diseases, we looked into three targets, namely DAO (red), PYGL (green), and VCP (blue), to understand the underlying market perception concerning patents. **B) Targets for insulin metabolism-related neurodegenerative diseases.** In the case of Alzheimer’s disease, we looked into a set of targets that play a role in insulin and sugar metabolism, namely RET (turquoise), INSR (orange), and SLC5A1 (violet). Each of the three targets follows a similar patenting profile. **C) Secretase-related targets in Alzheimer’s diseases.** The major secretase enzymes related to Alzheimer’s disease include ADAM17 (pink) and BACE1 (teal), and both of them follow a different patenting profile despite being in the same class. **D) Patenting profile for ELANE, a target within Alzheimer’s disease.**

Another illustrative target in the ranked list of targets was ATPase valosin-containing protein (VCP/p97). This protein plays a role in intracellular homeostasis by regulating protein metabolism and is associated with multiple diseases such as Paget disease and Amyotrophic Lateral Sclerosis and its types [32, 33]. However, due to the complex nature of the gene’s function in the body, research took an extended period, causing a dormant period in the patenting world. As soon as it gained interest, there was a steep increase in the number of patent documents, but this number soon fell **(Figure 9A)**. The key reason for this decline could be realistically attributed to VCP inhibitor CB-5083, a molecule developed by Cleave Bioscience, that failed at phase-1 due to off-target effects [34]. Subsequently, it was deduced that the covalent binding towards the target caused mutations that modified the morphology of protein, thus, making cells resistant to the drug [35]. Moreover, since the target is involved in cellular homeostasis [36], a deeper understanding of the MoA of drugs is needed for such targets.

Another target that surfaced in the rare disease domain with the highest number of associated patent documents was D-amino acid oxidase (DAO). It is a neuromodulator that was found to be involved in psychotic disorders [37]. This target has been active over the years from both the patenting **(Figure 9A)** and scientific literature, with several clinical trials. Additionally, the discovery of the target’s role in schizophrenia and cognitive dysfunction, and the need for its inhibition in these conditions, is one of the reasons for its active patenting landscape [38]. However, in the past two years, there has been a decrease in patents. This decline in patent activity could be ascribed to the COVID-19 pandemic, as global attention and resources have been focused on addressing the outbreak [39].

Interestingly, in the context of neurodegenerative diseases, some of the most well-known targets for Alzheimer’s, such as amyloid beta precursor protein (APP) and tau (MAPT) [40], were not found in the top target list **(Supplementary Table 2).** This was likely due to an absence of publicly available bioactivity data within resources such as ChEMBL on these targets, as antibodies and other biologicals are not considered. Contrary to these, coagulation factors, such as coagulation factor X (F10), which have a direct link to neuro-related diseases, were found in the top list [41] due to several novel small molecule modulators discovered. At the top of the target list was ret proto-oncogene (RET), with 886 patent application documents, demonstrating an active patenting profile **(Figure 9B)**. This activity profile can be attributed to the growing interest and research regarding insulin and sugar metabolism in the brain, where the receptor RET contributes towards metabolic homeostasis [42]. Other targets that contribute to the same mechanisms include insulin receptors (INSR) and glucose transporters solute carrier family 5 member 1 (SLC5A1), which were also found in the list later with an active patenting profile **(Figure 9B)**.

Secretases such as alpha-secretase (ADAM17) have gained interest within the pharmaceutical industry since the early 2000s due to their role in inhibiting amyloid beta formation [43]. Despite high patenting activity in the early years, there was a sharp decrease in 2005, and interest in such targets has been declining since then **(Figure 9C)**. Possible reasons for this decline include the failure of clinical trials for secretase inhibitors Lundbeck’s Flurizan and Lilly’s semagacestat in later years [44, 45]. The major drawback of these compounds was their inability to reach the desired concentration in the brain, as they could not pass the blood-brain barrier, thus, not reaching long residency time.

Other targets, such as neutrophil elastase (ELANE), also demonstrate patenting cliffs [25]. As described previously, this is correlated with either research activities on the target or clinical trial outcomes of drugs. The same pattern was observed in ELANE **(Figure 9D)**, which involved the development of elastase inhibitors and their advanced clinical trials [46]. The declining patenting activity was attributed to the identification of multiple mutations for the targets and the involvement of these mutants in multiple diseases. This target complexity thus required further research on both the biological and drug development sides.

Lastly, it is already known that there is a “lag” in the period between the filing of patent applications and the scientific research involved. We extrapolated this lag in the target space by looking into selected top targets and correlating the patent applications and research publications over time (**Supplementary Figure 4**). It can be observed that, on average, there is a lag of 5 years between the research involved on a target (with or without disease context) and the filing of patents relevant to the target. There are indeed certain outliers to this lag period such as INSR, which took at least 14 years to enter into the Alzheimer’s patenting world. One reason for this could be due to the importance of the target on other diseases, causing it to be extensively researched in the early 2000s. However, only after 2009 did the study of INSR in neurodegeneration start following the recognition of type 2 diabetes as a risk factor for Alzheimer’s [47].

### 3.4. Identification of drug repurposing scenarios of diseases using patent documents

As drug discovery is so resource intensive, a preliminary assessment of promising or under-exploited protein target and/or chemical scaffold combinations repurposing drugs could be a convenient alternative approach. We aimed to investigate whether drugs that have been repurposed for different therapeutic areas are also present in the patent corpora for the two disease areas. To achieve this, we clustered the patent documents based on the compounds they mentioned and ranked them in descending order of their counts. Since we were interested in repurposed compounds, we restricted the patent search to granted documents only. It is important to note that a patent application does not exclusively cover a single compound due to the presence of Markush structures [48]. These structures are a scaffold representation of a compound with the addition of variable side chains, thus making it more difficult for researchers to identify a specific compound via a direct search match, although commercial software tools (like SciFinder) have been developed to solve this problem. A single compound can also be covered by multiple patents, as the same compound can be used for varied innovative features (such as disease type or treatments, physical pharmaceutical forms, combinations with other therapeutic compounds, etc.), allowing for independent drafting for each area. Moreover, multiple patent drafting does not necessarily mean the compound is used for multiple purposes, as they may be patents filed for the compound’s formulations or non-obvious structural modifications. Thus, a systematic case-by-case study needs to be done to identify such compounds and ranking of compounds based on patent documents can serve as a starting point for such analysis.

Aligning our definition of repurposed compounds to a patent perspective, we categorised a compound as repurposed if it was found in more than two granted patent documents (belonging to B or E kind code). This resulted in a reduced dataset of 145 repurposed compounds in rare diseases and 1,928 in Alzheimer’s diseases, out of 585 and 22,682 compounds, respectively. Moreover, from the entire compound dataset, only four compounds for rare disease compounds and 215 compounds for Alzheimer’s diseases have been tested in clinical trials (as of 1st of Jan 2023 http://clinicaltrial.gov) (as shown in **Supplementary Figure 5)** with minimal overlap with the dataset of repurposed compounds. This indicates that most of the compounds within the granted patent space under study are still in the research or pre-clinical phase, likely due to the limited availability of bioassay data on clinically validated compounds in public data repositories such as ChEMBL.

Cleave Biosciences’s CB-5083 is one of the RD repurposed drugs forming an interesting case study. The company had patented the compound in Europe (EP-2875018-B1) and the United States of America (US-10010554-B2) as an inhibitor of VCP for cancer treatment. Following a phase I clinical trial, which revealed off-target activities, clinical development was halted [49]. Despite the unsuccessful clinical trial, it has made its way to the RD community regarding its potential use in treating Paget disease, a VPC-dependent disease [32]. Another example is Fosdagrocorat, a compound patented by Pfizer as a glucocorticoid receptor modulator for osteoporosis (EP-2114888-B1). Research has linked the same receptor to rheumatoid arthritis [50], thus making it possible to repurpose the compound for this rare disease [51]. Upadacitinib, commonly known by the trade name Rinvoq, is an Abbvie-developed kinase inhibitor that has been approved by the FDA for the treatment of multiple diseases, such as rheumatoid arthritis [52] and atopic dermatitis. It inhibits the Spleen tyrosine kinase (Syk) (US-10072034-B2), which has recently been identified as a key modulator of tauopathy, a disorder involving the accumulation of Tau [53]. This discovery opens up the potential for the drug to be used to treat neurodegenerative diseases. Similar to rare diseases, we found a few compounds derived from the cancer domain. Sotorasib is a compound that inhibits a specific KRAS mutant, KRAS G12C and demonstrates anti-cancerous properties (US-10519146-B2) [54]. Several researchers have identified several KRAS mutants that play a role in brain and spinal-related haemorrhage [55]. This presents an opportunity to test the compound on multiple KRAS mutations and optimise it to create a mutant-specific or general-purpose compound. In summary, it is evident through the above examples that the mining of patent documents from a compound perspective can provide insights into how compounds that have been repurposed for multiple disorders beyond the scope of their original patents.

### 3.5. Limitations and future perspective of the approach

The current study illuminates the historical trends observed in neurodegenerative and rare disease drug discovery and research through the analysis of patents. However, it is important to note that the methodology employed has limitations due to the lack of inclusion of “biologicals” compounds such as vaccines or long peptides. One such limitation is the indirect relationship between proteins and patents, as the connections were made based on chemical modulators, and the initial proteins may not necessarily be mentioned in the patent. To address this, further analyses using natural language processing (NLP) methods to extract gene annotations in patent documents are the best alternative. Additionally, the modulators linked to the proteins were sourced from open-source data repositories, such as ChEMBL. As the deposition of clinical candidate structures is infrequent and often only occurs after early clinical trials, the patent analysis may miss important compounds currently in development. To mitigate this, we focused on highly active compounds with submicromolar activity to increase the chances of identifying “champion” compounds in a patent. Lastly, the time range chosen for the analysis (2000-2021) may have skewed the overall distribution of results, as certain compounds approved for treatment before 2000 were not included in the analysis. For instance, Jeffrey Cummings (2018) mentioned in his article that “Agents producing cognitive enhancement may have mechanisms independent of AD-specific pathology (e.g., 5-HT6 antagonists)” [56]. 5-hydroxytryptamine receptor 6 (5-HT6) antagonists like memantine and acetylcholinesterase inhibitors have been approved for AD treatment and only alleviate some AD symptoms by enhancing cholinergic signalling. These chemicals are not curative and do not surface in our analysis, as they were patented before 2000. Despite these limitations, the study aims to provide insight into the intellectual effort required within the pharmaceutical industry to identify competitors’ patents containing “clinical candidates” and establish a successful “me-too” strategy.

## 4. Conclusion

Finding safe and effective therapies for diseases with unmet patient needs is a crucial goal of pharmaceutical research. Traditionally, the discovery process was initiated by systematically reviewing the scientific literature for the most promising disease-linked therapeutic targets or pathways. Once a targeting mechanism has been selected, it would either be subsequently validated or eventually discarded and an alternative strategy adopted based on new experimental insights or literature analysis. Additionally, assessment of target opportunities and estimating their potential for successful translation in development candidates is a core strategic activity within the pharmaceutical industry. Such analysis is driven by machine learning models trained on large multimodular data, including toxicological effects and druggability characteristics. Moreover, the novelty associated with the chemical structure of candidate molecules is a critical aspect of patent assessment, as companies need to be aware of pre-existing successful scaffolds. This chemoinformatic integration of structural features is often a key factor in defining medicinal chemistry strategies when optimising leads and further progressing compounds to development stages.

This study uses the PEMT tool to extract, analyse, and explore the patent landscape for rare and Alzheimer’s diseases. The results revealed historical trends in the past 20 years and identified key patent holders and application dates. Such a retrospective trend paved the way for generating a longitudinal visualisation of the importance of the targets in both diseases. Additionally, the analysis showed the potential for using patent literature in drug repurposing and annotating the clinical significance of targets. Last, but not least, the study was able to identify and reflect the shift between the research and patenting period involved in drug discovery. Certainly, this effort cannot be assumed to reach the same proficiency as a professional patent analysis as it takes advantage of public databases, which themselves are limited, as discussed in the previous sections. This paper is focused on the integration of public bioactivity datasets with patent applications and their analysis. We will inspect through future analyses how the patent information can optimise drug search within disease mechanisms networks, which is out of the scope of the current paper. This is just the tip of the iceberg as huge effort and resources need to be dedicated to standardising the data and making them a potential source for future ML and LM models.

Our work demonstrates that the PEMT tool provides a comprehensive and informative patent cohort to understand the research in the field of small molecule drugs. This landscape not only aligns with scientific literature but also allows for competitive intelligence analysis in pharmaceutical R&D. Overall, this patent landscaping approach from a drug discovery perspective highlights the importance of analysing patents and their usefulness in making decisions for future drug development efforts. The possibility of tracking patenting activities from publicly available resources might help define strategies for target ranking within a therapeutic focus. Our methodology will contribute towards simplifying and streamlining the process of target prioritisation in drug discovery campaigns at a pharmaceutical company.

## Author contribution

AZ and YG conceived the work. YG programmed PEMT. YG and AZ performed the analysis and contributed to ideation. AZ and YG have written the manuscript. All the authors contributed towards reviewing the manuscript. All authors have read and approved the final manuscript.

## Acknowledgements

The authors thank Daniel Domingo-Fernández, Bruce Schultz, and the anonymous reviewers for their comments and suggestions on the manuscript. We would also like to thank Bruce Schultz for providing the protein data for the HBP network, hosted by Fraunhofer SCAI.

## Funding

This work was supported by the German Federal Ministry of Education and Research (BMBF) [01GM2003A] (project ‘TreatKCNQ’).

## Notes

https://github.com/Fraunhofer-ITMP/Pharmaceutical-patent-landscaping

### Competing Interest Statement

The authors have declared no competing interest.

### Summary of Updates

Figure and legend information for Figures 3 and 6 have been updated.

