## Supplementary data for "Pharmaceutical patent landscaping: A novel approach to understand patents from the drug discovery perspective"

#### Supplementary Figure

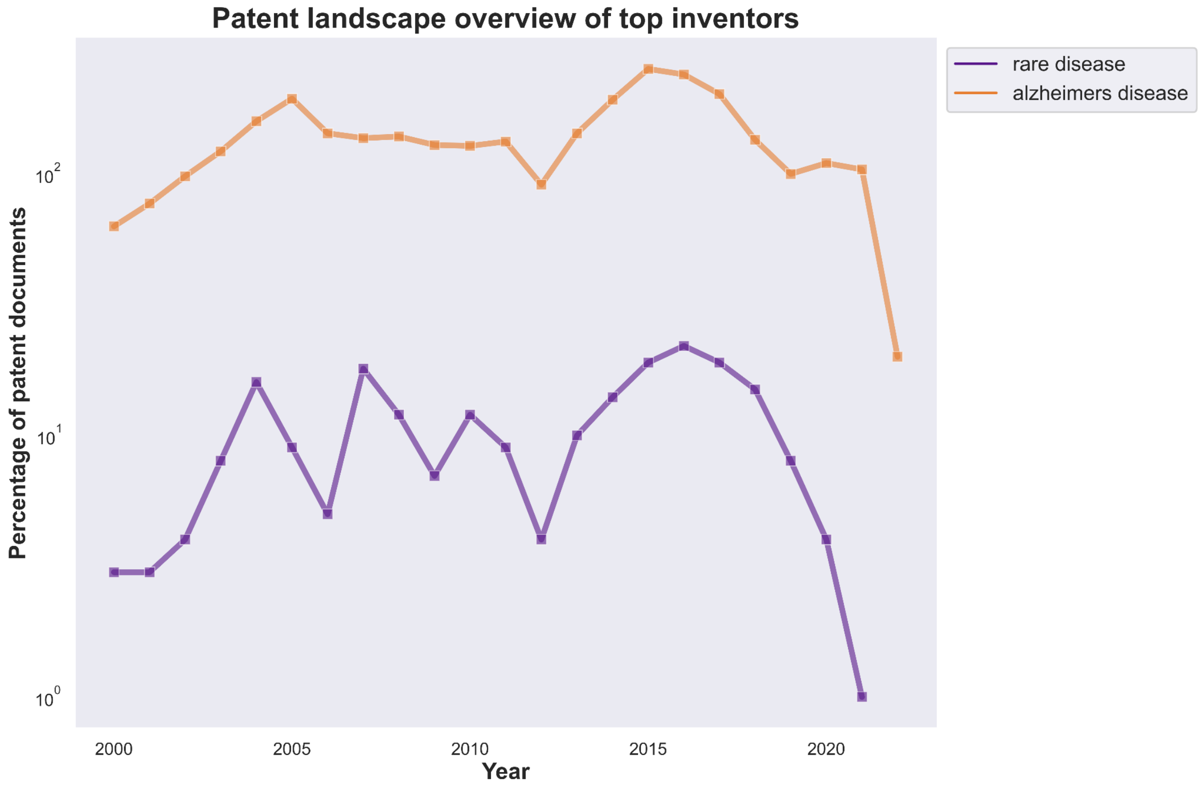


**Supplementary Figure 1. Overall patenting activity across the top assignees in the different disease indication areas**. The figure indicates the cumulative patenting activity growth of top 10 assignees within each indication area. It can be seen that compared to rare diseases, the Alzheimer disease's patenting activity is 10 times more. Also, an evident drop within the patenting activity for the two indication areas can be obsered during the start of pandemic around 2021.

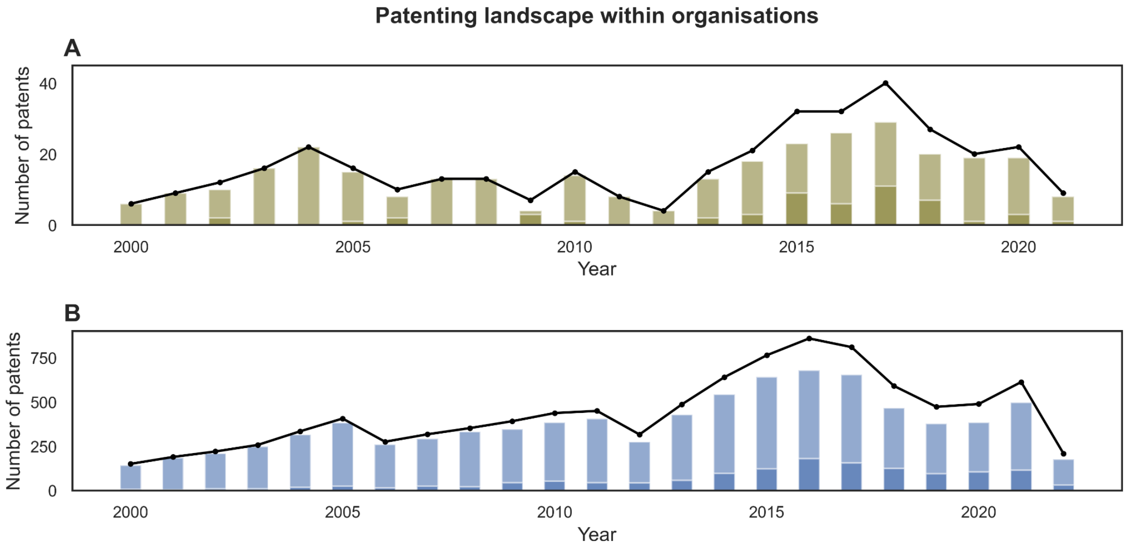


Supplementary Figure 2. Time-dependent patenting activity with organisation subclasses. An exponential increase in patenting can be observed along with a decrease during the COVID-19 period (2020-21). The annual patenting activity for each year can be observed by the line plot on top of the bar plots. A) The patenting landscape between the industry and academic sector for rare diseases where the darker green bar indicate the patenting activity for university while the light green indicate the activity by industry. B) The patenting landscape between the industry and academic sector for Alzheimer’s diseases where the dark blue indicates patenting activity by universities and light blue the activity by industry.

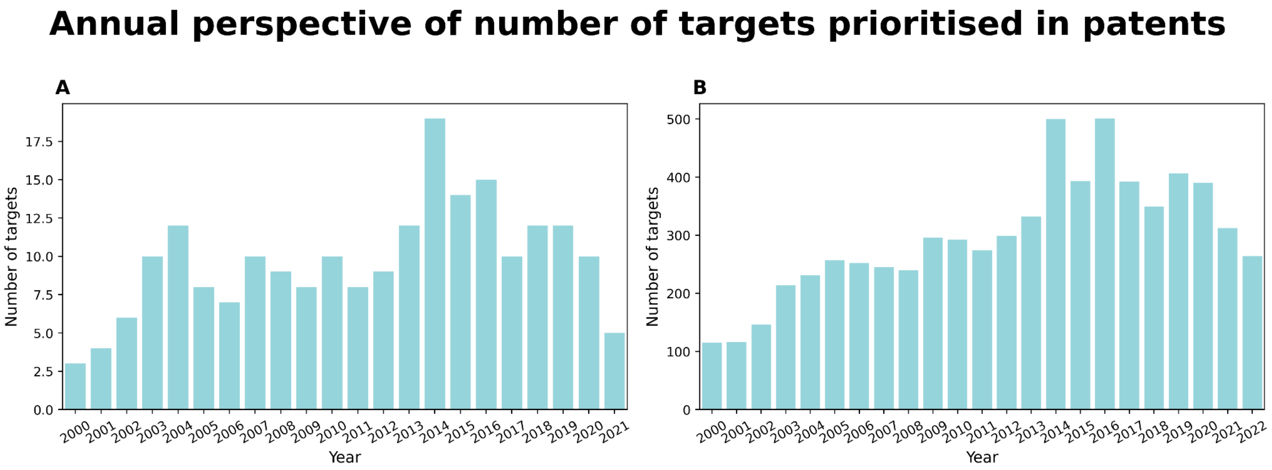


**Supplementary Figure 3. Plot indicating the annual perspective of disease targets in (A) rare diseases** and **(B) Alzheimer’s disease**. The bar plots show the distribution of the number of targets that were considered to be patented per year. The cliffs and valleys in both plots indicate that the underlying target space changed over the years.

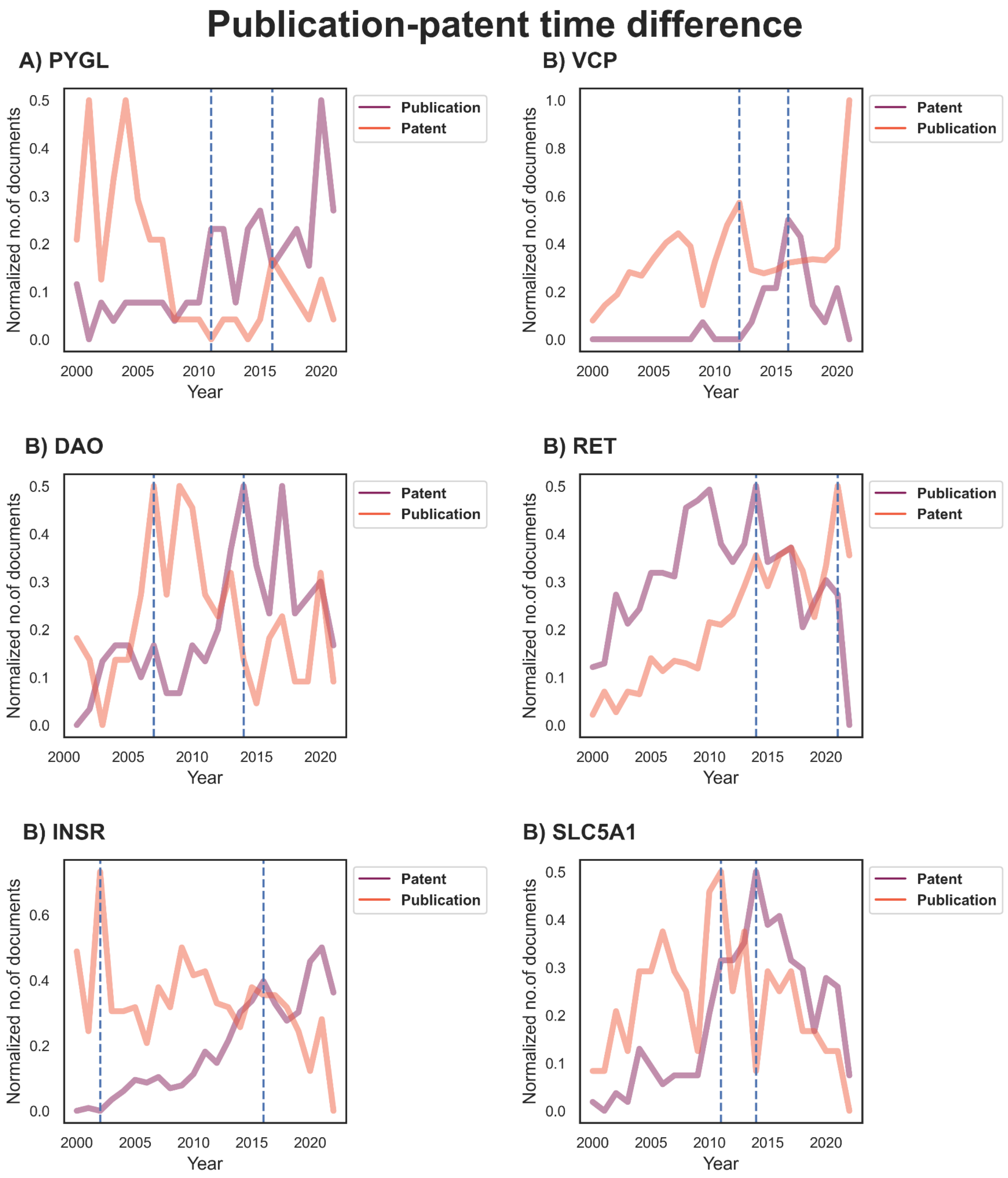


**Supplementary Figure 4. Time lag between research and patents.** The distribution plot denotes the difference in the time period between filing patents for a target and research involved around the target. The x-axis denotes the year, while the y-axis denotes the normalised document (research paper or publication) count for the given year. In each plot, two dotted lines are drawn that indicate the first peak observed in the publication or research area and the peak observed in the patenting area (post the publication peak) in a sequential manner. Only top targets based on the patenting documents are shown here as examples.

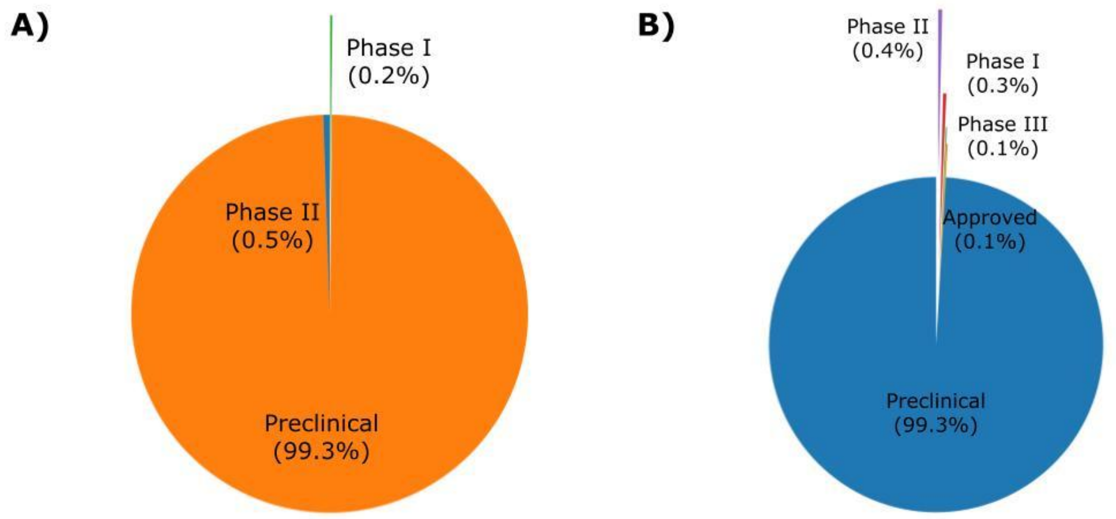


**Supplementary Figure 5. Chemical clinical phases within granted patent documents. A)** The compounds in rare diseases found in granted patents belong to either preclinical, phase I, or phase II. **B)** The compounds in neurodegenerative diseases found in granted patents belong to phase III and phase IV or approved drugs in addition to the classes found in rare disease compounds.

### Supplementary Tables

| HGNC target symbol | Name of target | Number of patent documents |
|---|---|---|
| DAO | D-amino acid oxidase | 127 |
| STS | Steroid sulfatase | 115 |
| PYGL | Glycogen phosphorylase L | 71 |
| PTPN2 | Protein tyrosine phosphatase non-receptor type 2 | 39 |
| CYP19A1 | Cytochrome P450 family 19 subfamily A member 1 | 28 |
| VCP | Valosin containing protein | 27 |
| GABRD | Gamma-aminobutyric acid type A receptor subunit delta | 23 |
| SLC9A3 | Solute carrier family 9 member A3 | 19 |
| ASAH1 | N-acylsphingosine amidohydrolase 1 | 17 |
| NOD2 | Nucleotide binding oligomerization domain containing 2 | 17 |

**Supplementary Table 1. Ranked top targets for rare diseases.** The table provides an overview of the total number of patent documents associated with the top 10 targets in the field of rare diseases.

| HGNC target symbol | Name of target | Number of patent documents |
|---|---|---|
| RET | Ret proto-oncogene | 886 |
| CYP2D6 | Cytochrome P450 family 2 subfamily D member 6 | 747 |
| PLK4 | Polo like kinase 4 | 662 |
| CSF1R | Colony stimulating factor 1 receptor | 595 |
| MET | MET proto-oncogene, receptor tyrosine kinase | 574 |
| SLK | STE20 like kinase | 553 |

| | | |
|---|---|---|
| MAP4K4 | Mitogen-activated protein kinase kinase kinase kinase 4 | 548 |
| FER | FER tyrosine kinase | 544 |
| F10 | Coagulation factor X | 535 |
| PTK2B | Protein tyrosine kinase 2 beta | 516 |

**Supplementary Table 2. Ranked top targets for Alzheimer's diseases.** The table provides an overview of the total number of patent documents associated with the top 10 targets in the field of Alzheimer's diseases.
